# Df-1 Cell Adaptation and Immune Responses of Lc-75 Vaccinal Strain of *Infectious Bursal Disease Virus* in Chicken

**DOI:** 10.1101/2024.07.09.602647

**Authors:** Fentahun Mitku Abate, Destaw Asfaw Ali, Belayneh Getachew, Hawa Mohammed

**Affiliations:** College of veterinary medicine, Tuskegee university, Alabama, USA; Department of Veterinary Paraclinical Studies, College of Veterinary Medicine and Animal Sciences, University of Gondar, Gondar, Ethiopia; Research and Development Department, National Veterinary Institute, Bishoftu, Ethiopia

**Author notes:** Correspondents need to be addressed to: Fentahun Mitku Abate.

**Keywords:** Adaptation, Chicken, *DF-1 cells*, Infectious bursal disease, Immunogenicity, Immunosuppression, *Ethiopia*

## Abstract

The current *Infectious bursal disease virus* (*IBDV*) vaccine strain of LC-75 in Ethiopia is being produced through chicken embryo fibro blast cell which has short lifespan and limited need of specific pathogen free eggs. Ideally, vaccine should have better longevity and being effective in cost and time. Experimental research was conducted to adopt DF-1 cell and to validate the immune responses, immunosuppressive effect and immunogenicity tests, of LC-75 vaccinal strain of *Infections bursal diseases virus* A total of 76 chickens were used for these experiments. The seroconversion rate of the vaccine were measured using haemaglutination inhibition (HI) test in the vaccinated two experiment groups, adapted IBDV and Newcastle vaccines at two weeks interval for the first group and only Newcastle vaccine for the second experiment groups. To perform the immunogenicity test, two groups having 20 chickens per group were used and samples of serum were taken on 0-, 7-, 14- and 28-days post vaccinations and subjected to indirect ELISA test. The group that received both vaccine types produced haemagglutination inhibition titer (HIT) of 97.33±22.49 whereas the other group that received only Newcastle vaccine produced 124±24.92. The control group always showed no detectable antibody while the vaccinated group was able to produce average antibody S/P values of 0.00 ± 0.01 at day 0, 0.02± 0.01 at day 7, 1.05±0.10 at day 14 and 0.83±0.05 at day 28. RT-PCR using a 400 bp IBDV viral protein 2(VP2) specific primer resulted from positive bands in all samples. In conclusion, the vaccinal strain was able to replicate and adapt on the DF-1 cells and it was found to be immunogenic as well as less immunosuppressive.

## 1. INTRODUCTION

Chicken farming makes a significant contribution to the provision of high-quality food in the form of meat and eggs for young children in Sub-Saharan Africa’s well-balanced diet [1,2]. However, insufficient feed inputs, poor management, infectious illnesses, and a lack of appropriate selection and breeding practices continue to limit poultry production’s contribution to small-holder farmers and the country’s economy [2, 3]

In Ethiopia, poultry farms are expanding at a faster rate than ever before [4]. As poultry farming became more intensive, epidemics of newly acquired diseases began to emerge [5, 6]. Infectious bursal disease (IBD) is one of the illnesses that is posing a threat to chicken production rates and product quality. Despite frequent vaccination protocols and better biosecurity measures, the illness has become a critical problem in commercial and backyard chicken production systems. Furthermore, most control measures developed in the country do not account for indigenous chickens, which may result in most strategies failing [3]. Since 2004, mutation and genetic reassortment have resulted in the emergence of highly pathogenic strains [7]. As a result, immunization is thought to be a crucial way to safeguard hens during their early weeks of life [8].

Due to its highly contagious acute viral illness that affects chickens aged 3-6 months, infectious bursal disease (IBD) is a top priority disease [9]. IBD is caused by the serotype 1 *Infectious Bursal Disease Virus* from the genus Avibirnavirus and family Birnaviridae [10]. It is a major chicken immunosuppressive virus that can exacerbate earlier infections with other infectious agents and limit the bird’s ability to respond to immunization by causing humoral and cellular immune responses to be damaged [11, 12]. The virus prefers lymphoid tissue, the bursa, the thymus, the spleen, and the bone marrow. As a result of the virus’s destruction of lymphocytes and macrophages, it affects the immune system, causing vaccination failures and concomitant infections in the poultry farming [8, 13, 14]. The molecular characterization and pathogenicity of the *vvIBDV* (serotype 1) from natural infection of turkey could also possibly spread to chickens causing severe economic losses [15].

Live vaccines for *IBDV* have been developed and are classified as mild, moderate, or hot depending on their virulence. Mild vaccinations are safe for specific pathogen free hens, but they are ineffective in the face of significant maternal antibodies or against IBDV strains that are extremely virulent. Although intermediate and hot vaccines are far more efficacious, they can cause moderate to severe bursa fabricius lesions [16]. For the prevention of IBD, a variety of vaccinations with varying levels of protection are available. The most widely used vaccinations are live attenuated vaccines, inactivated oil-emulsion adjuvant vaccines, and recombinant IBDV-vp2 protein vaccines [17, 18]. Some oral administration of the mixed herbal extract for 5 weeks can stimulate the immune response to *IBDV* vaccination and relieves the pathogenicity of *IBDV* co-infection in chickens [19].

Ethiopia has been using chicken embryo fibroblast cell to produce live IBDV vaccine strain LC-75. Because these are primordial cells, they take a long time to produce, demand a lot of labor, and require the importation of specified pathogen free eggs. They also have a short lifespan. The vaccine’s production costs will be decreased if it is adapted in DF-1 cell lines, because DF-1 cells are continuous cell lines that can be easily produced in a short period of time. Under normal culture conditions, these cells are excellent as substrates for virus propagation, recombinant protein expression, and have a high rate of proliferation [20]. The objective of this work was to develop a DF-1 cell line-based *IBDV* vaccine manufacturing platform using the LC-75 vaccinal strain, as well as to assess the strain’s immunogenicity and immunosuppressive effects on chicken.

## 2. MATERIALS AND METHODS

### 2.1. Experiment Site

The experiment was conducted at National Veterinary Institute (NVI) in Bishoftu, Ethiopia. The institute is a government-run entity that develops and distributes veterinary vaccines around the world. This company holds various international accreditation certificates (ISO/IEC 17025:2005) in vaccine manufacture and illness diagnosis from international accrediting bodies (NVI public communication office).

### 2.2. Study Animals and Design

One day old white leghorn chickens were used as study subjects. The chicken were obtained by hatching from eggs of breeder flocks at the NVI. The parent flock was raised in NVI under strict hygiene and biosecurity conditions [21]. The chickens were specific antibody negative for IBDV. For the course of the trial, feed and water were provided ad libitum. At the end of each set immune response experiments (immunosuppression and immunogenicity experiments separately), the chicken were euthanized humane way using cervical dislocation technique. After adaptation of the vaccine on the DF-1 cells, two experimental designs were applied, one for immunosuppressive effect and the other for the immunogenicity test. For these two experiments a total of 76 chickens were used; 36 for the immunosuppressive effect test and 40 for the immunogenicity test.

#### Experiment one: Immunosuppressive testing of the adapted virus

The IBD vaccination adapted from DF-1 cells was given as an eye drop. Each of the 12 chicken was inoculated with the supernatant (0.2 ml/chicken). As controls, two other groups of 12 chicken of the same age were housed separately. Each bird in the IBDV-vaccinated group and one of the control groups received one field dose of thermostable live ND vaccine at the age of two weeks. After two weeks (14 days) of ND vaccine administration, the haemagglutination inhibition (HI) response of each chicken was evaluated, and protection against challenge with a virulent strain of ND virus was measured. The severity of the *NDV* challenge was validated using the second control group, which was retained without IBDV or ND vaccination. If the HI response and protection provided by the ND vaccination is significantly lower in the group given both vaccines than in the group given only Newcastle vaccine, the IBD vaccine is considered immunosuppressive [22].

#### Experiment two: Immunogenicity tests of the DF cell adapted virus

For the immunogenicity tests the DF-1 cell adapted vaccine was administered to chicken in ocular route as per the recommendation [21]. After administration of the vaccine, the level of the humoral response was measured serologically using indirect ELISA test. To do this, two groups of chicken containing 20 chickens per group were used. Each of the chicken in the treatment group was administered a supernatant of 0.2ml of the adapted vaccine and the other group was left as uninoculated control. Finally, serum was collected at days; 0, 7, 14, and 28 and tested using ELISA. The level of the antibody induced against the administered vaccine was measured in terms of the serum to positive control ratio (S/P ratio) from the ELISA reader connected with software package.

### 2.3. Study Materials

#### 2.3.1. Media and solutions used

According to the manufacturer’s instructions (HIMEDIA Laboratories, India), growth medium and maintenance medium were made from Glasgow’s minimal essential medium (GMEM) with 10% and 2% sterile fetal calf serum (FCS), respectively. Another media and solutions such as Trypsin versine solution and Phosphate buffered saline (PBS) were utilized as needed.

#### 2.3.2. Experimental viruses and DF-1 cell lines

The working master seed of *Infectious bursal disease vaccine* strain (LC-75), Df-1 cell, Newcastle challenge virus and thermostable Newcastle vaccine were obtained from NVI different departments.

### 2.4. Study Techniques

#### 2.4.1. Passage of adherent monolayers of DF-1 cells

The DF-1 cell seed was first resurrected from liquid nitrogen and cultivated in a 75cm^2^ cell culture flask using cell medium. After that, it was placed in a CO_2_ incubator and observed under an inverted microscope (Olympus CK2, Japan) to see if a complete monolayer had formed. The DF-1 cells were then transferred to a class-II safety cabinet after being confirmed confluent (Lab care, England). To remove dead cells, the medium was taken and washed three times with pre-warmed PBS. A pre-warmed 0.25% trypsin solution was used to suspend the confluent monolayer. The suspended cell in the flask was then separated into three 25cm^2^ tissue culture flasks using GMEM and cultured at 37°C in a CO_2_incubator with 5% CO_2_. Under the inverted microscope, the cells were inspected twice daily for the creation of a full monolayer [23, 24]

#### 2.4.2. Inoculation of IBDV (LC-75) to DF-1 cells

Virus infection was performed on healthy and confluent monolayers of DF-1 cells 36 hours after sub-culturing. With 70% ethanol, the working space under the laminar air flow cabinet was sterilized. The maintenance media and PBS were warmed in a 37°C water bath. In a 25cm2 flask containing a confluent monolayer, the growth media was removed and the cell monolayer surface was washed three times with pre-warmed PBS. After that, the DF-1 cells were infected with 1 ml of *IBDV* (LC-75) with a titer of 6.4 EID50/ml and incubated at 37°C in a CO_2_ incubator. For all passage stages, a control flask of fresh cells with confluent mono layers was preserved under comparable conditions. From the time of inoculation until six days later, the infected cells were studied twice a day under an inverted microscope for the production of CPEs. Virus-infected cells were extracted six days after inoculation, labeled as passage 1(P1), and kept at −20°C overnight. The P1 virus was then frozen thawed three times and injected onto a new monolayer of DF-1 cells and CPEs were monitored twice a day for the next six days. The virus was also extracted six days after infection, labeled as P2, and stored at −20°C overnight. P3 virus was obtained by a third infection, and CPEs were examined twice a day for the first six days after inoculation.

#### 2.4.3. Infectivity assay of the adapted virus

Each passage’s viral suspension was diluted in sterile tubes from 10^-1^ to 10^-5^ by combining 0.5ml viral suspension with 4.5ml GMEM. Secondly, 100 μl DF-1 cells were dispensed into 96 micro plate wells in a way to have ten replicates for each dilution. Then 100μlsuspension from each of the dilutions was added to all the wells containing Df-1 cells (from columns A to E in the 10 rows). For negative controls, column 11 was left empty and column 12 was vaccinated only once. Finally, the plate was sealed with a microplate sealer and incubated for 6 days at 37°C with 5% CO_2_. Starting on the day of inoculation and continuing until day six, the inoculated plates were examined twice daily under an inverted microscope. The titers for each virus passages were determined according to the spearman formula. Log_10_ 50% end point dilution = (x_0_ − d/2 + d ∑ r_i_/n_i_)); where x_0_ = log_10_ of the reciprocal of the highest dilution (lowest concentration) at which all cell monolayers are positive; d = log_10_ of the dilution factor(the difference between the log dilution intervals);∑ r_i_/n_i_) = sum proportion of the tests beginning at the lowest dilution showing 100% positive results;r_i_ = number of positive monolayers out of n_i_[24, 2].

Enzyme-linked immunosorbent assay (ELISA) was used to assess antibody titers to *IBDV*. The ELISA technique was used in accordance with the commercial Infectious Bursal Disease Antibody indirect ELISA diagnostic Kit’s instructions (ID.VET innovative diagnostic, France). By comparing the value of the unknown to the positive control mean, the presence or absence of IBDV antibody was identified. The antibody level was calculated in terms of S/P ratios using the algorithm provided in the ELISA kit. Serum samples with S/P ratios less than or equal to 0.3 were classified as negative, whereas those with S/P ratios more than 0.3 were positive.

To determine the extent of interference of the adapted IBDV vaccine from ND antibody production, the HIT of the treated groups was measured and compared. A HI test employing *Newcastle disease virus* antigen was used to check the sera for particular antibodies that inhibited haemaglutination. Equal volumes [25] of eight haemaglutination units of the NDV were mixed with serial twofold dilutions of serum in PBS. 25 liters of cleaned chicken red blood cells were added and incubated for 30 minutes at room temperature. The reciprocal of the last dilution of serum that entirely prevented haemaglutination was used to calculate the titer. The test was carried out in accordance with the national veterinary institute’s test procedure standard of operation (NVI-VPS-RD-PR-06). For Newcastle disease and avian influenza, the HI test is regarded a gold standard for assessing the humoral immune response [15,22].

#### 2.4.4. The validity test and interpretation was conducted using ELISA

The test sample was considered as negative if the S/P ratio is less than 0.3 or the antibody titer is less than 875, otherwise positive. According to the kit, test validity was also examined, only if the positive control mean optical density value (ODpc) is greater than 0.025 and the ratio of the positive and negative controls mean values is greater than 3 were valid (ID.VET innovative diagnostic, France).

#### 2.4.4. Molecular identification of DF-1 cell adapted LC-75 vaccine

Reverse transcriptase polymerase chain reaction (RT-PCR) was conducted using to the molecular laboratory protocol (NVI-VPS-RD-PR-05). All three passage levels were subjected to molecular identifications in order to confirm the adopted virus. The virus infected cells supernatant fluid was collected for RNA extraction. The QiagenRNeasymini Kit technique was used to extract RNA (Cat.nos.2010210). RNase-free water was used to elute the RNA and kept at −20°C. The isolated RNA was then used as a template to make a 25μl master mix. RNase free water (4μl), 10mMdNTPs (1μl), PCR buffer x (5μl), Q solution 5x (5μl), each primer (2μl), one step RT-PCR enzyme mix (1μl), and RNA template (5μl) were all included in the mastermix. For this reaction, forward pri mer 5’TCTTGGGTATGTGAGGCTTG 3’and reverse primer 5’CCCGGATTATGTCTTTGA 3’ were used (IBDV-P2-PANVAC). The samples were loaded into the thermocycler (Applied Biosystems 2720). Then the machine was adjusted at 50°C for 30 minutes for DNA synthesis (1cycle), at 95°C for 15 minutes (for 1 cycle) for initial denaturation followed by 35 cycles of second denaturation at 95°C for 30sec, at 55°C for 30sec for annealing, at 72°C for 30sec for elongation and finally followed by one cycle of final elongation at 72°C for 7 minutes (NVI-VPS-RD-PR-09).The PCR product was analyzed by gel electrophoresis. A 10μl sample of the PCR reaction mixture was electrophoresed and separated on a 1.5% agarose gel in Tris EDTA buffer. After staining with gel red (0.5µg/ml), the electrophoresis was run for 1:20 hour at 120V and PCR products was visualized by viewing the gel with an ultraviolet light and photographed and compared with a100 bp DNA ladder (marker). The result of gel picture was captured by the camera, saved and printed out and documented (NVI-VPS-RD-PR-42) [24].

### 2.5. Data Management and Statistical Analysis

The cultured flasks were observed daily for any morphological changes and the observed change was recorded accordingly. The serological data was entered to Micro Soft Excel sheet program to be analyzed using STATA version 14 and subjected to independent t-test. Descriptive statistics was also employed to summarize the data. Statements of statistical significance was decided based on the when alpha 0.05%.

## 3. RESULTS

### 3.1. Results of the Adaptation Process

In this experiment the LC-75 vaccinal strain was inoculated to a monolayer of DF-1 cells up to three serial passages, all of which showed signs of CPEs such as rounding, detachment from the bottom of the culture flask and cellular aggregations when observed under the inverted microscope and the cell lines were susceptible to IBDV. The cultured flasks were observed on a daily basis but cytopathic effects were started to be seen from the third days of infection in passage one and from second day of both passage two and three. At the 5^th^ and 6^th^ days of post infections, the CPE’s were typical. But the control uninoculated flasks did not show any cytopathic effects, there was normal cellular morphology ofDF-1 cells (Figure 1). Upon titration (expressed in log10 TCID50) all the three passages produced average titers of 10^5^, 10^5.5^and 10^6^TCID50/respectively.

**Figure 1.**
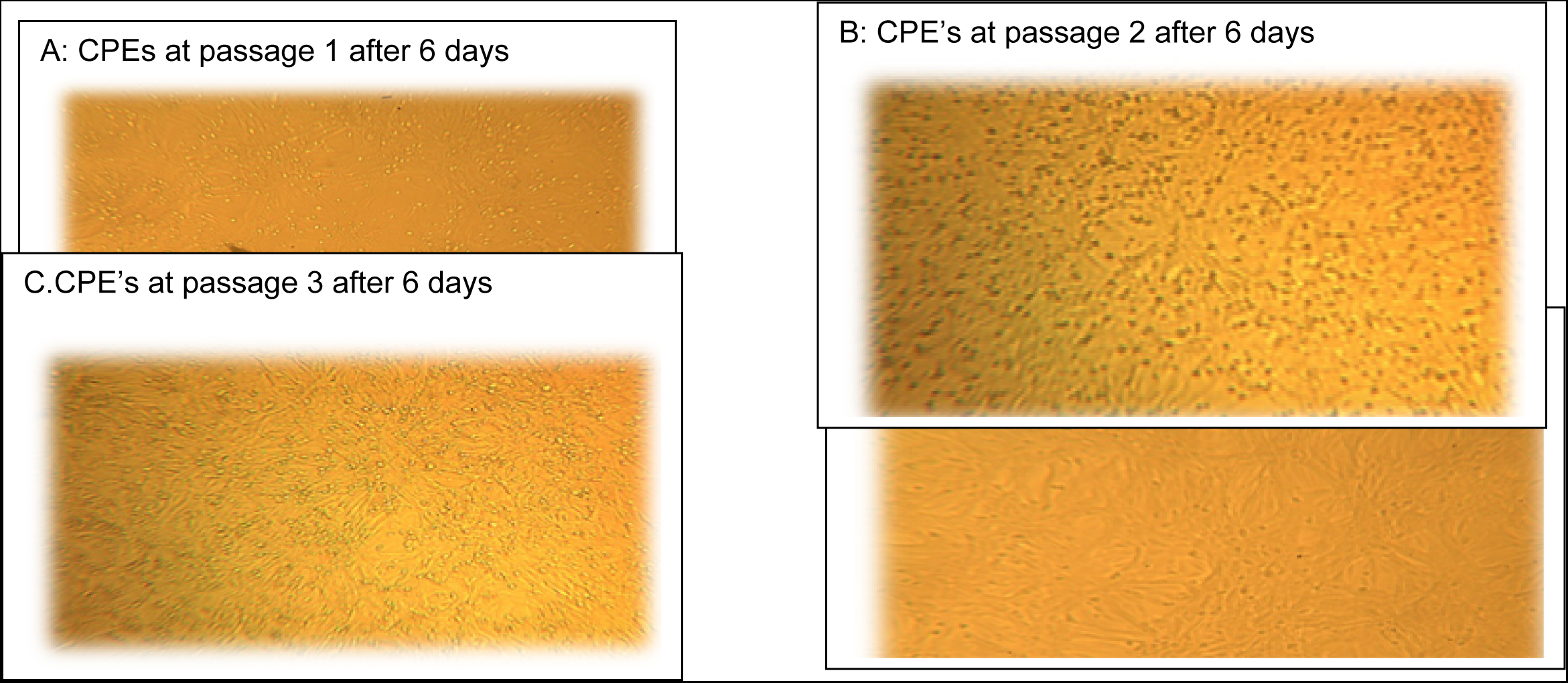
(A-D). LC-75 IBDV vaccine strain grown on DF-1 cell

### 3.2. Results of the Immunosuppressive Effect Test

The non-vaccinated chicken showed clinical signs of new castle starting the second day of post challenge such as respiratory distress (gasping), dragging of the feathers to the ground and greenish diarrhea. Death was started after the fourth day of post challenge. After nine days of post challenge nine out of twelve chicken died, only three of them survived, but they showed mild clinical signs. However, chicken from the two treatment groups (group 1 & 2), showed no overt clinical signs of Newcastle (Table 1).

**Table 1.** Clinical observations noticed after challenging by virulent NCD virus strain.

| Groups | No. of chicken per group | Death after challenge | Survived | Morbidity after challenge | Mortality after challenge |
| --- | --- | --- | --- | --- | --- |
| **G1(IBD & NC vaccinated)** | 12 | 0 | 12 | 0(0%) | 0(0%) |
| **G2(Newcastle vaccinated)** | 12 | 0 | 12 | 0(0%) | 0(0%) |
| **G3(unvaccinated)** | 12 | 9 | 3 | 12(100%) | 9(75%) |

To see whether there was a difference in seroconversion ability to the Newcastle vaccine in the two vaccinated groups (between group 1 and 2), the HI titer results were measured and compared. The HI titer in both groups was positive as well as protective (all sera showed HI titer>16). Group 1(received IBDV & ND vaccines) produce an average HI titer of 97.33±22.49 and group 2(received only Newcastle vaccine) produce an average HI titer of 124± 24.92. From this result there is a slight difference of the average HI titer but it’s not statistically significant (P>0.05) (Table 2).

**Table 2.**
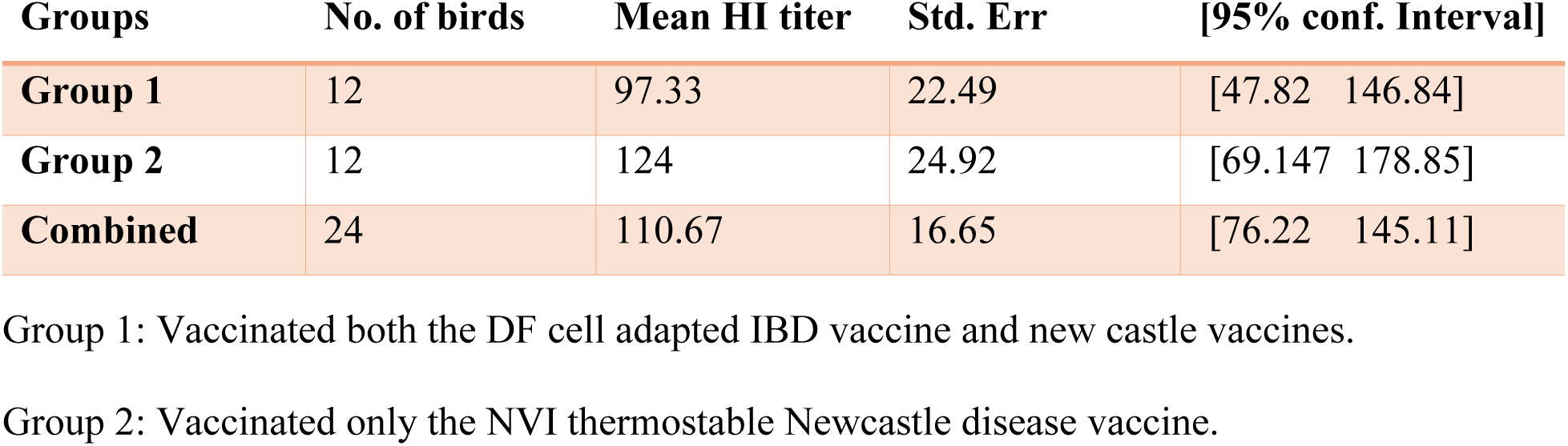
Mean serum HI antibody titer of the chickens in the two treatment groups.

### 3.3. Results of the Immunogenicity Test

To see the level of the humoral immune response of the experimental chicken, the sera S/P values of the vaccinated and the control groups was measured and compared. All the non-vaccinated chicken had antibody S/P ratio much less than 0.3 and all the vaccinated ones produced antibody level having S/P ratio much higher than 0.3. Thus, no meaningful serological comparison could be made between the two groups. The antibody level produced at days 0, 7, 14, and 28 of post vaccination in the vaccinated group was compared. The average S/P values were: 0.00 ± 0.06 at day 0, 0.06± 0.01 at day 7, 1.05±0.10 at day 14 and 0.83±0.05at day 28. There is a slight decrease of S/P values from 14 days to 28 days, but that difference was not significant (p> 0.05). The average serum antibody levels (S/P ratio values) (Table 3).

**Table 3.**
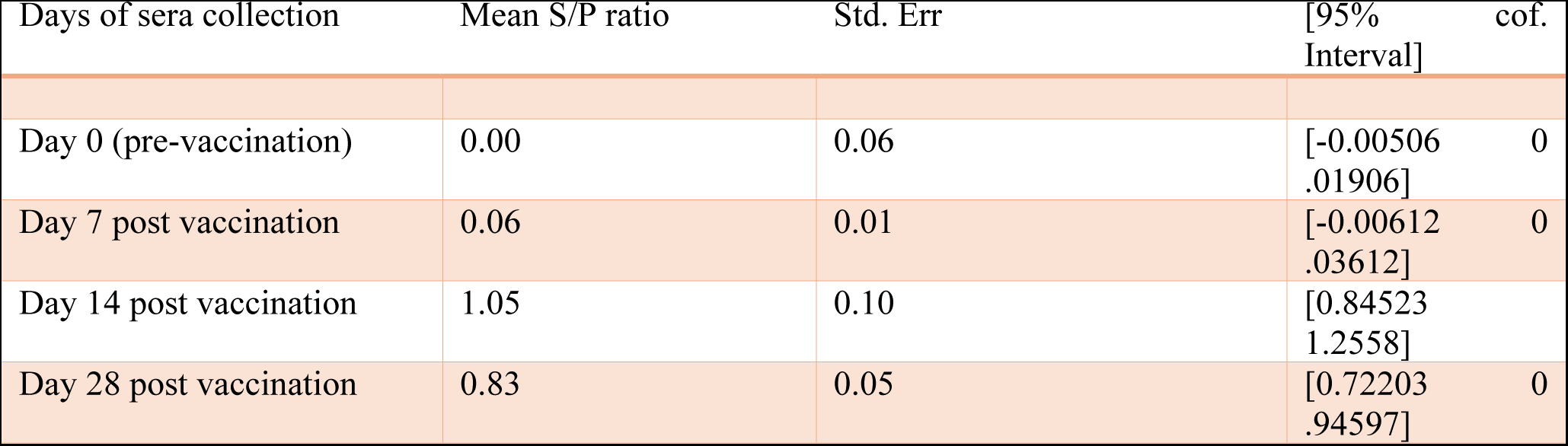
Mean serum antibody sample to positive (S/P) ratios in the vaccinated group.

Cut off values provided on the Kit: S/P ≤ 0.3 is considered as negative (not protective) whereas S/p> 0.3 is positive and protective. According to these cutoff values indicated in the Indirect ELISA Kit for IBDV (ID.VET innovative diagnostic, France), the antibody level induced at pre-vaccination (at day 0) and after seven days of post vaccination was very low so that it could not protect the chickens from natural infections or challenge by virulent IBDV. However, the antibody levels produced from all the experimental chickens at 14th and 28th days of post vaccination are protective from IBDV virus infections or challenge. It reached at its peak at 14^th^ day and slightly decreased at 28^th^day of post vaccination. This pattern of the antibody induction is shown graphically in the figure below (Figure 2).

**Figure 2:**
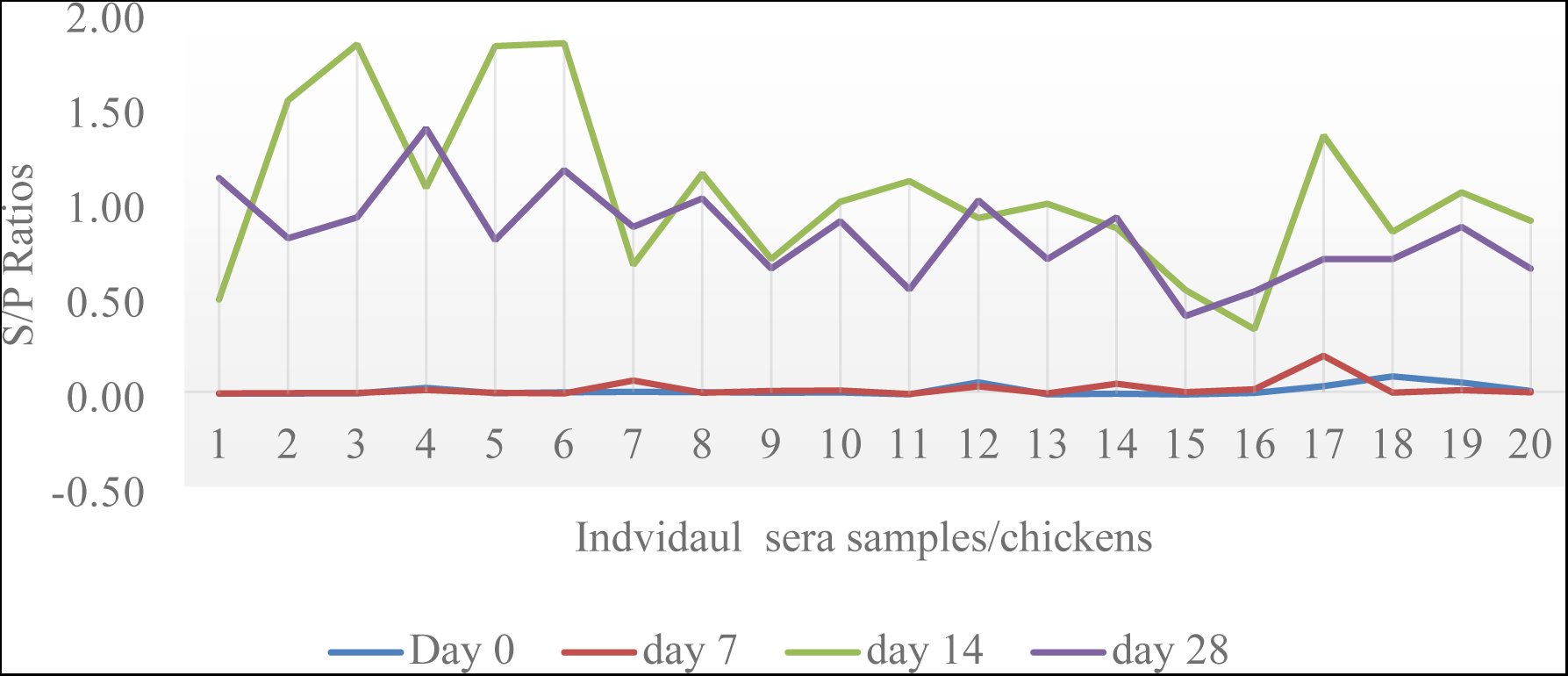
Graphical representations of the S/P ratios during the experimental periods

### 3.4. Molecular Identification Test Results

The identification process was done in duplication of each passage level. RT-PCR using a 400 bp IBDV viral protein 2(VP2) specific primer resulted positive bands in all the six samples. All the six samples were aligned exactly with the band having 400 base pairs (Figure 3)

**Figure 3.**
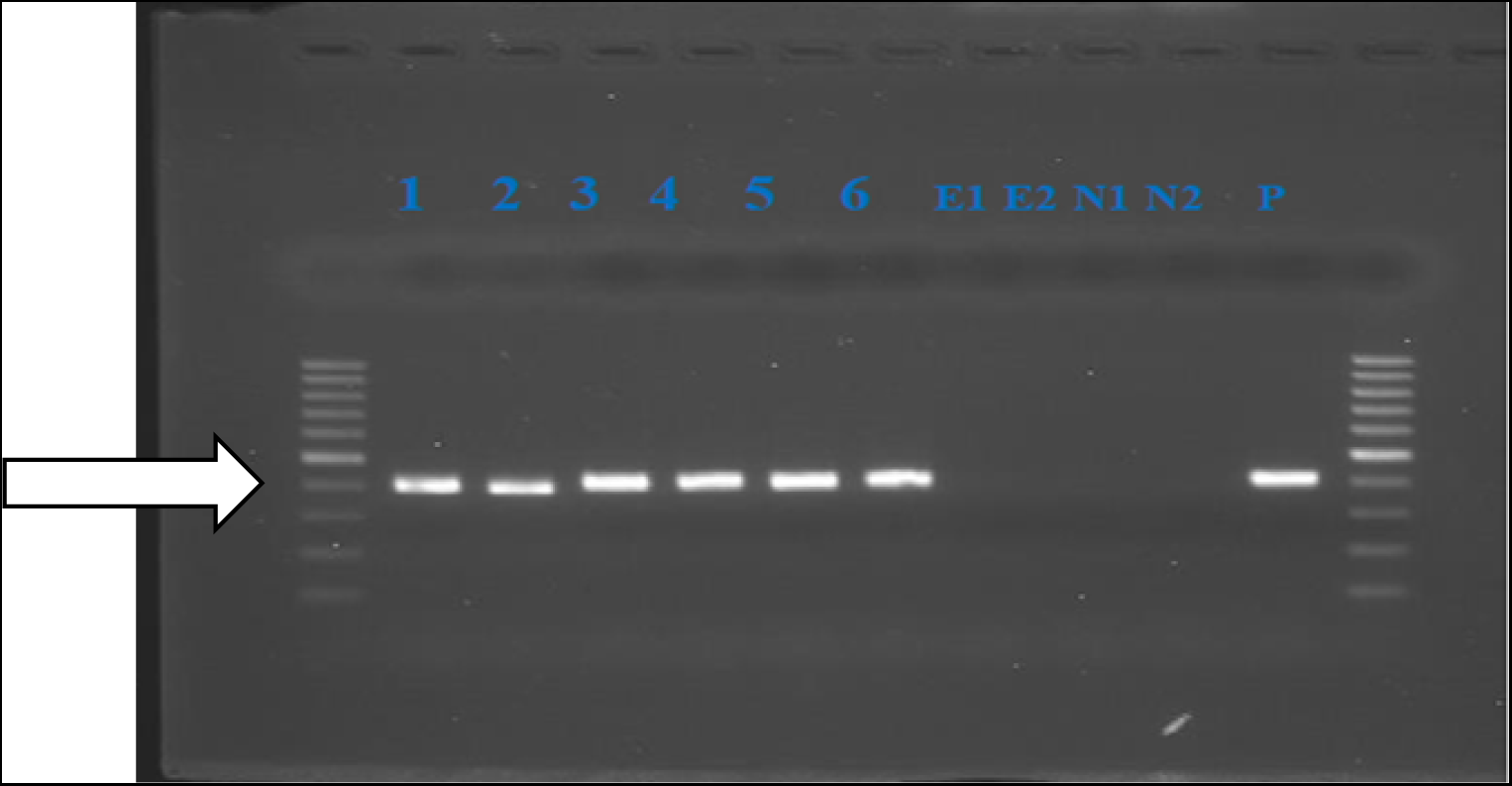
Molecular results captured after gel electrophoresis of the samples. M= 100bp molecular markers; Lanes 1-6 =sample from the 3 passage levels (2 from each); E1 and E2 =Extraction control; N1 and N2= negative controls and P= positive control

## 4. DISCUSSION

*Infectious bursal disease virus* is susceptible in a variety of mammalian continuous cell lines. Infectious bursal disease virus (IBDV)-susceptible DF-1 cell lines have also been shown to exhibit cytopathic effects in response to viral infection [26]. IBD vaccinal strain infections on DF-1 cells caused cytopathic effects characterized by cell aggregation, cell rounding, fragmentation of cells into small particles, and finally detachment from the substrate, until the monolayer was destructed, according to the results of this study’s adaptation trial. The cytopathic effects were detected at all levels of passage. This finding is comparable to that DF-1 cells are susceptible to infection by creating pronounced cytopathic effects (CPEs). This cell line has a better growth potential and infectivity titer, both of which are beneficial in vaccine production [27]. The adapted vaccine’s RT-PCR output exhibited accurate amplification of a 400bp amplicon specific to the primers for the hypervariable, VP2 gene fragment, indicating that the vaccine virus was successfully replicated and adapted on these DF-cell lines while keeping its original molecular identity. This finding is consistent with Hamoud who used primers with the same amplification capacity as VP2 gene segments to detect IBDV from formalin-fixed paraffin-embedded tissue [28, 29].

The immunosuppressive impact of the adapted vaccinal strain is one of the experiments covered in this work. Other researchers have previously showed immunosuppression in hens that have been infected with Infectious Bursal Disease at a young age. Acute illness and death are thought to occur during outbreaks as a result of the virus’s necrotizing action on the tissue of the Fabricius bursa in chicken [30]. If the birds survive and recover from this stage of the disease, they will stay immunocompromised, making them more susceptible to other infections and making vaccines against other viruses ineffective. The virulent strain vaccinations of IBDV, according to Hair-Bejo (27) may induce severe bursal lesions comparable to those seen in field epidemics of the virus. Virulent vaccines have also been observed to cause subclinical illnesses in birds, resulting in decreased performance [31].

Research conducted elsewhere are in support that virulent IBDV such as LC-75 have immunosuppressive effects of variable degrees. However, the findings of this test results appear to support that the DF-1 cell adapted Vaccinal strain of LC-75 has lees deleterious effect on the bursa of vaccinated birds since the average NDV-HI titers produced between the two groups was not significantly different. In addition both the vaccinated groups (the one that received both IBD and Newcastle) and that received Newcastle vaccine alone, were equally protected by the virulent Newcastle challenge virus. There was no any morbidity and mortality observed during the ten days of post challenge. However, the non-vaccinated groups showed 100 morbidity and 75% mortality. The findings of this test result is in line with the findings of the same work done by Sadrzadeh[32] who found that there were no significant differences of NDV-HI titers among groups received both IBD and Newcastle vaccines or Newcastle vaccine alone. This result is also in agreement with the findings of Kulikova [33] who indicated that LC-75 strain induced lower bursal damage and better performance parameters.

A vaccine candidate must elicit a significant level of antibody titer and this antibody titer must be of adequate length to be successful [34]. In the immunogenicity test of this study, the adapted virus vaccine produced a high level of antibody (expressed in terms of S/P ratio) which is in agreement with [35] This antibody level was highest at the 14 days of post vaccination but was extremely low at day 0 and 7 of post vaccination. This time of onset of the antibody is in accordance with Jakka[36] who reported low(non-protective) antibody level at 7 days of post vaccination in India, but slightly different with the findings of El-Bagoury[37] in Egypt who reported that protective level of antibody was observed stating to the first week of post vaccination using an oil adjuvant IBDV vaccine. This difference might be associated with the dose and the formulation, and the preparations and/or the co-administration of the adjuvant in the previous study. Another source of variation might be due to the age and the type of management of the experimental birds.

After fourteen days of post vaccination, the immune response was at a peak level (average S/P ratio of 1.05±0.10). This figure is comparatively agreed with the result of Mekuriaw [21] who found an antibody level with average S/P ratio of 0.558 at fourteen days post of vaccination but less than the present study. This difference might be occurred due to the breeds. The former used bowman brown breeds in their experiment, opposed to white leghorn breeds in the present study. The other difference might be associated with the presence of residual maternally derived antibody (MDA) in their study chicken since they were bought from farms where vaccination of the parental flock against IBDV might be took place regularly.

Individual chickens in the vaccinated group were all able to produce antibody levels with S/P ratios greater than 0.3. According to the ELISA Kit, a high level of antibody protects chickens from IBDV-induced illnesses. Despite the lack of a challenge test in our study, the antibody level appears to be consistent with the findings of Hossain [38] and Otsyina and Aning[38], who both reported that chickens vaccinated with live virulent vaccines were protected when challenged ten days later with highly virulent IBDV vaccine strains. Zaheer and Saeed [34] found that outbreaks of IBD in flocks inoculated with a range of vaccine strains, which contradicted these findings. Other triggering factors, such as the immunological status of chicks at the time of vaccination, inadequate handling and delivery of vaccines, or inappropriate vaccination programs, appear to have an impact on the success of IBDV immunization [39]. The average antibody level began to fall slightly four weeks after vaccination (after 28 days), although this was not statistically significant when compared to the level produced two weeks after vaccination (after 14 days). This minor decrease in antibody levels may indicate the necessity for revaccination to achieve excellent protection against bursal disease virus infection in a flock [39].

## CONCLUSION

This work presented that the LC-75 Vaccinal strain is well adapted and replicated in the DF-1 cell lines. The adaptation was confirmed by the presence of typical cytopathic effects and molecular identifications. The cells were able to produce a virus with substantial average titer/ml which warrants the use DF-1 cells for replication of the LC-75 vaccinal strain of *IBDV* for production of large scale and high quality *IBDV* vaccine. The adapted vaccine humoral immune responses revealed that the vaccination could produce a high antibody titer, which might protect the chicken against IBD. The immunosuppressive potential of the adapted vaccine was also found to have less deleterious effects on the chicken. The time it takes to maintain its protective level and efficacy of the adapted vaccine should be evaluated in terms of a challenge virulent *Infectious bursal disease virus*.

## Author Contributions

Conceived and designed the experiments: FM, DS and BG. Performed the experiments: FM, BG and HM. Analyzed the data: FM and DS. Contributed reagents/materials/analysis tools: BG and HM. Wrote the paper: FM and DS

## Acknowledgments

We express our sincere gratitude to the NVI management members, particularly Dr. Esayas Gelaye and Dr. Takle Abayneh, for their unwavering facilitation and all forms of assistance related to the project. We also deeply appreciate the cooperative and steadfast technical support provided by the technician staff at the National Veterinary Institute throughout all the laboratory tasks.

## Notes

### Competing Interest Statement

The authors have declared no competing interest.

